# Exploring the space of human exploration using Entropy Mastermind

**DOI:** 10.1101/540666

**Authors:** Eric Schulz, Lara Bertram, Matthias Hofer, Jonathan D. Nelson

## Abstract

What drives people’s exploration in complex scenarios where they have to actively acquire information? How do people adapt their selection of queries to the environment? We explore these questions using Entropy Mastermind, a novel variant of the Mastermind code-breaking game, in which participants have to guess a secret code by making useful queries. Participants solved games more efficiently if the entropy of the game environment was low; moreover, people adapted their initial queries to the scenario they were in. We also investigated whether it would be possible to predict participants’ queries within the generalized Sharma-Mittal information-theoretic framework. Although predicting individual queries was difficult, the modeling framework offered important insights on human behavior. Entropy Mastermind opens up rich possibilities for modeling and behavioral research.

## Introduction

Humans are curious animals. From learning how to speak to launching rockets into space, exploration drives mankind’s progress small and large. Human exploration has been studied in self-directed learning paradigms in adults and children, in domains including causal learning (Bramley, Dayan, Griffiths, & Lagnado, 2017), categorization (Meder & Nelson, 2012), control (Osman & Speekenbrink, 2012), and explore-exploit tasks (Wu, Schulz, Speekenbrink, Nelson, & Meder, 2018). Some experiments have used games including Battleship (Gureckis & Markant, 2009) and 20 questions (Nelson, Divjak, Gudmundsdottir, Martignon, & Meder, 2014). Self-directed learning can lead to improved performance (Gureckis & Markant, 2012; Markant, Ruggeri, Gureckis, & Xu, 2016). For instance, participants actively intervening on a causal system made better inferences about the underlying causal structure than subjects who received identical information passively (Lagnado & Sloman, 2004).

Recent conceptual work (Coenen, Nelson, & Gureckis, 2018; Gureckis & Markant, 2012; Schulz & Gershman, 2019) is underpinned by the assumption that behavior is goal-directed and that people select observations based on a metric of usefulness (Settles, 2009). What metric best predicts how people evaluate the usefulness of possible queries? Past work has focused on the expected reduction of uncertainty, the extent of predictions’ improvement, or the maximization of future rewards (Nelson, 2005). One study optimized experimental materials to maximally distinguish between different measures in an experience-based probabilistic classification task (Nelson, McKenzie, Cottrell, & Sejnowski, 2010). Results showed that participants were better described by probability gain than by information gain or other measures.

Markant and Gureckis (2012) tested whether participants maximize payoffs or information gain in a game of “battleships” (Gureckis & Markant, 2009), where each query cost money and an attempt to maximize utility would lead to different queries than information-gain based strategies. Surprisingly, participants’ sampling behavior was nonetheless best matched by information gain. The authors argued that using information gain would lead to more knowledge about the underlying structure and therefore can be an effective strategy, no matter what the final task will be. Similar results have been obtained in an active causal learning task (Bramley, Lagnado, & Speekenbrink, 2015).

Exploiting the characteristics of the Entropy Mastermind game, we investigate people’s sensitivity to the information structure of their environment (mathematical entropy or psychological uncertainty) and adaptive strategy selection when facing different levels of probabilistic uncertainty. In particular, we focus on what information metrics best predict how people evaluate the usefulness of possible queries, and on what initial-guess strategies people use.

## A quintessential game of exploration

In the Mastermind code-breaking game, both information search and exploitation are essential for breaking the code. Thus, Mastermind offers a potential platform for bringing together pure information models (like expected information gain) and reinforcement learning models. In the classic two-player version of the game one player generates a secret colour code (e.g. blue, red, green) and the other player has to guess the secret code by repeatedly testing codes (making queries) and receiving feedback about the correctness of items in the guessed code. Although Mastermind has been extensively studied in computer science (for references see Berghman, Goossens, & Leus, 2009; Knuth, 1976), comparatively less work has been done in cognitive science (but see Laughlin, Lange, & Adamopoulos, 1982; Zhao, van de Pol, Raijmakers, & Szymanik, 2018).

We introduce the game “Entropy Mastermind” for studying exploration-driven problem solving and uncertainty reduction (Fig. 1). Key attributes of Entropy Mastermind, which distinguishes it from the classic game, are that Entropy Mastermind is a single-player app-based game in which hidden codes are drawn from known, and typically nonuniform, probability distributions. The probability distribution from which the hidden fruit code is drawn is depicted as a “fruit bowl” icon array. The player is informed that the items are mixed before each draw, and drawn with replacement to form the hidden fruit code. Thus, Entropy Mastermind makes it possible to research how the level of entropy affects people’s strategies and efficiency in game play.

**Figure 1:**
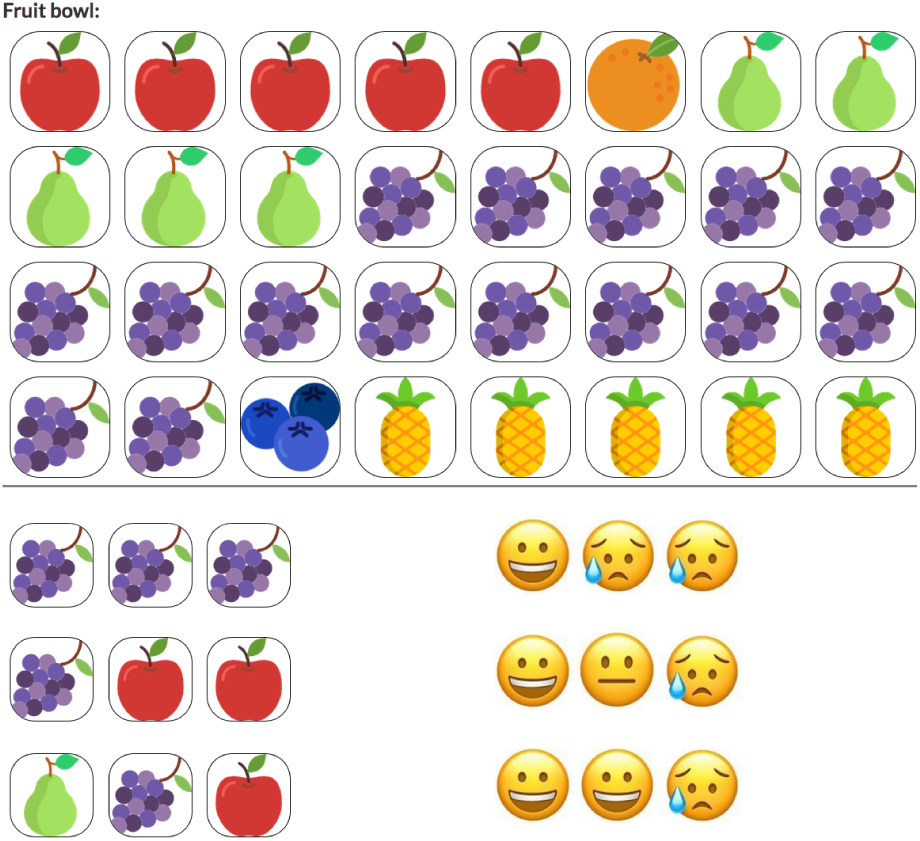
Fruit Salad Mastermind: High Entropy Condition. **Top:** Icon array presenting an example fruit bowl that generated the secret code. Probability distributions follow one of four entropy recipes, resulting in low, medium low, medium high and high entropy levels. Fruit types are apples, oranges, blueberries, grapes, pears, and pineapples for all possible versions of the fruit bowl. Codes are generated by randomly sampling fruits with replacement. Duplicates are allowed, so it is possible that the same fruit could appear in all positions of the hidden code. Players have to guess which fruit is in which position of the three slots of the secret code, by clicking on the position they want to change. Each position is initially blank; clicking cycles through the possible fruits. Once participants are satisfied with the proposed code, they can click on a “Check” button (not shown), and then receive feedback. **Bottom:** History of game play illustrating feedback. In the first guess, the player guessed 3 grape items. The feedback (one smiling face followed by two frowning faces) conveys that exactly one of the items is correct in type of fruit and in location. However, the player does not know which of their guesses is correct. There is no correspondence between the position of the guess and the position of the feedback: happy faces always come first, then neutral and lastly frowning faces. In the second guess, the player tested grape in the first position, and apple in each of the other two positions. The feedback (smiling face, neutral face, frowning face) indicates that one of the items is the correct type of fruit in the correct location, another item is in the code but needs to be moved to a new location, and another item is not in the code at all. As before, the guesser has to figure out which feedback face corresponds to which item in the code. The third guess of pear, grape, apple obtains two smiling faces and one frowning face. At this point the guesser can infer that the middle position is grape, and the final position is apple; the guesser must still figure out the first item.

As a first step toward modeling behavior in a probabilistic framework, we use a model that values both maximizing the probability of a correct query and a curiosity bonus, similar to recent work on human reinforcement learning (Schulz, Konstantinidis, & Speekenbrink, 2018; Wu et al., 2018). The curiosity bonus can be defined as information gain in the space of possible hypotheses (hidden codes). Whereas information gain has traditionally been thought of as reduction in Shannon entropy, any entropy metric could be used. We use the Sharma-Mittal space of entropy measures (Sharma & Mittal, 1977), which provides a framework within which many different kinds of entropy measures arise. According to the setting of two parameters, known as the *order* and *degree*, this entropy space can recover Shannon entropy, Bayes’s error, and entropies from the Arimoto, Rényi and Tsallis families of entropy measures, among others (Crupi, Nelson, Meder, Cevolani, & Tentori, 2018). One of our research questions is whether Entropy Mastermind can help identify which model of uncertainty best predicts exploratory behavior.

In what follows, we formally define the Sharma-Mittal space as a unifying framework for information gain measures. We then report a preliminary study assessing and modeling human behavior in Fruit Salad Mastermind, a version of Entropy Mastermind in which the code jar is a fruit bowl and items are different kinds of fruits. First results show that participants adapted their queries to the level of entropy in the environment, solving games in less entropic environments more efficiently than in more entropic environments. Thus, basic assumptions for using Entropy Mastermind as a model of an information environment varying in entropy were met. Both the exploration and exploitation parts of the model were important to account for human behavior. However, distinguishing between different parts of the Sharma-Mittal space turned out to be difficult. Future research could work towards designing tasks that are optimized for the purpose of discriminating among specific entropy models.

## Mapping the space of exploration

In Mastermind both *learning* about the true code and *guessing* the true code are important. To make this intuitive, suppose that there are two possible codes, given everything that has been learned to date, and that one of these codes has 90% probability of being the correct code. The same information, namely which code is correct, will be gleaned from testing either code; thus, the queries have equal value irrespective of which model of information gain is used. But clearly it is sensible to test the code that has 90% probability of being correct, thus having 90%, rather than 10%, probability of ending the game after the next query. We implement this idea via a softmax response rule on a value function which is based on the probability of each query being the correct code in the immediate time step, as well as a curiosity-driven exploration bonus^1^:

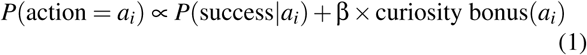

How promising a code seems is determined by its current probability of being correct *P*(*success*|*a*_*i*_). This probability is always the same given a specific history of queries and feedback. The curiosity bonus(*a*_*i*_) is weighted by a free parameter β and can be defined as how much an action promises to reduce uncertainty over the space of possible hypotheses (i.e., how much it reduces uncertainty about possible codes).

The uncertainty in a discrete random variable *K* = *k*_1_, *k*_2_, … *k*_*n*_ can be measured by its entropy. We use the generalized Sharma-Mittal space of entropy measures, that unifies multiple past proposals (Crupi et al., 2018), and can be defined as:

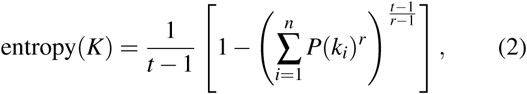

where *r* is the *order* and *t* the *degree* of the entropy measure. Note that limits, which exist, are used for points where the above equation is undefined. Although the above equation may not be immediately intuitive, there are a number of ways to build understanding about this space. All of the Sharma-Mittal entropy measures can be thought of as quantifying the average surprise that would be experienced if the value of the random variable *K* was learned. In the case of Mastermind, this is the average surprise that would be experienced if one were to immediately learn the true hidden code.

The degree parameter *t* governs which kind of surprise is averaged. If *t* = 1, then *surprise*(*k*_*i*_) = *ln*(1/*P*(*k*_*i*_)), as in Shannon and all of the Rényi entropies. If *t* = 2, then *surprise*(*k*_*i*_) = 1 − *P*(*k*_*i*_), as in the cases of Quadratic entropy and Bayes’s error. If *t* > 1, a test is more useful if it is conclusive than if it is not. If *t* < 1, a test is always less useful if it is conclusive than if it is not. The order parameter *r* determines what kind of averaging function is used. It can be thought of as an index of the imbalance of the entropy function, which indicates how much the entropy measure discounts minor (low probability) hypotheses. For example, when *r* = 0, entropy becomes an increasing function of the mere number of the possible options. When *r* goes to infinity, entropy becomes a decreasing function of the probability of a single most likely hypothesis. For further discussion and examples see (Crupi et al., 2018).

Several special cases exist within the Sharma-Mittal space, as Figure 2 illustrates. For example, Shannon entropy is the result of setting *r* = *t* = 1, and probability gain (also called error entropy) is the result of setting *t* = 2 and letting *r* → ∞. One of the goals of the present research is to investigate whether people’s striving for information (the curiosity goal) can be represented well as a generalized information gain metric, where information is defined as the expected reduction in one of the Sharma-Mittal entropy functions over the probability distribution of the possible codes.

**Figure 2:**
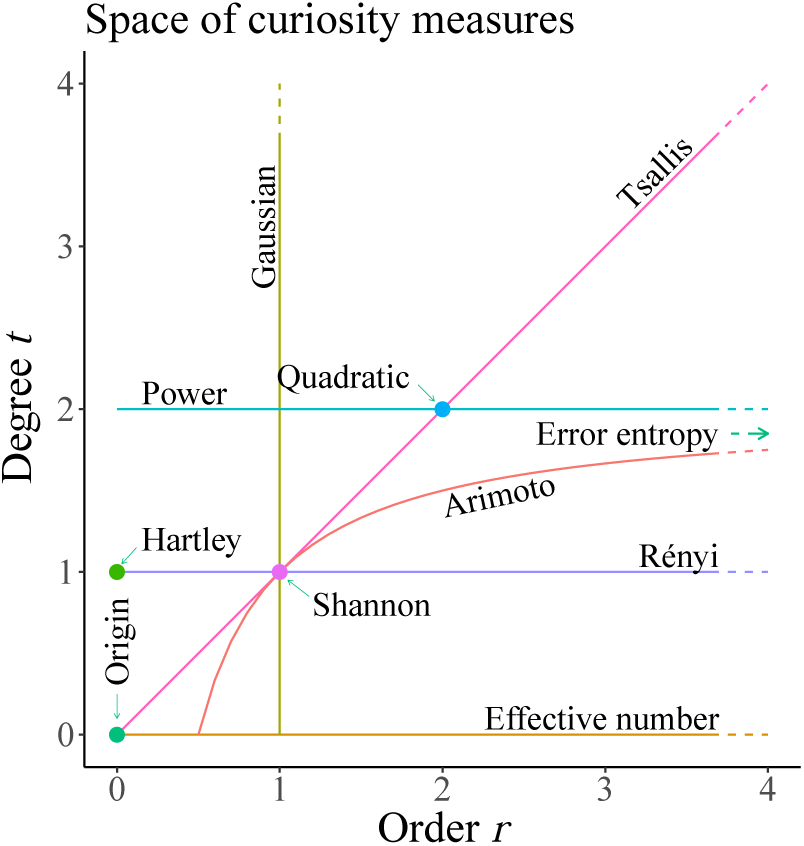
Sharma-Mittal space. The Sharma-Mittal family of entropy measures is represented in a Cartesian quadrant with values of the order parameter *r* and of the degree parameter *t*. The order parameter captures how much minor hypotheses are disregarded (e.g. that grapes may be contained in the code when the fruit bowl contains only a small proportion of grapes) and the degree parameter captures how prominent the goal of getting as close as possible to the state of certainty is (i.e. how much one strives to falsify existing hypotheses, e.g. that grapes are contained at all in the code). Each point in the quadrant corresponds to a specific entropy measure, each line corresponds to a distinct one-parameter generalized entropy function. Several special cases are highlighted.

## Methods

### Participants and Design

Forty-seven first-year undergraduate students (38 female, *M*_*age*_=19.04; *SD*=1.04; range: 18 to 23) at University of Surrey participated in our study as part of a cognitive psychology class. Participants gave informed consent in accordance with the University’s procedures and the Helsinki Declaration. They were introduced to the rules and interface of the game and completed a pretest. Participants then played Fruit Salad Mastermind, spending an average of 10.5 minutes on the task.

### Materials and Procedure

Participants were required to correctly answer four comprehension questions before game play began. These questions tested participants’ understanding of the goal of the game and the interpretation of feedback (i.e. making sure that they understood that the position of the faces did not correspond to the position of items in the entered code). Participants were instructed to figure out the secret code using as few guesses as possible. Since the experiment was self-paced, the number of rounds played varied between participants.

### Entropy conditions

In each game, one of the four entropy conditions was chosen at random and the six fruits were randomly assigned to the six proportions of that condition. The resulting generating “fruit bowl” was presented to participants as an icon array above the current game. A “hidden fruit code” was generated from that distribution. In the *very high entropy* condition, the secret code was sampled based on the proportions (5, 5, 5, 5, 6, 6). This means, for example, that there could be 5 pineapples, 5 apples, 5 pears, 5 blueberries, 6 grapes, and 6 oranges, out of a total of 32 items, from which three fruits were sampled with replacement to generate the secret code. In the *high entropy* condition, the secret code was sampled based on the proportions (1, 1, 5, 5, 5, 15). In the *low entropy* condition, the secret code was sampled based on the proportions (1, 1, 1, 4, 4, 21). Finally, in the *very low entropy* condition, the secret code was sampled based on the proportions (1, 1, 1, 1, 1, 27).

### Behavioral results

We analyzed behavioral results using both frequentist and Bayesian statistics. For testing hypotheses regarding the behavioral data and the model comparison, we used the default two-sided Bayesian *t*-test for independent samples with a Jeffreys-Zellner-Siow prior with its scale set to 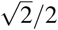 (Rouder, Speckman, Sun, Morey, & Iverson, 2009).

We first analyzed the number of required guesses to solve a game as a function of the entropy condition (Fig. 3a). This revealed a positive average rank correlation between how much entropy a condition contained and the number of queries participants required to solve a game (Kendall’s τ = 0.48, *t*(46) = 12.44, *d* = 1.81, *BF* > 100). More specifically, participants required fewer queries on average for the very low entropy games as compared to low entropy games (*t*(46) = −5.69, *p* < .001, *d* = 0.83, *BF* > 100). They also required fewer queries for the low entropy games than for the high entropy games (*t*(46) = −3.16, *p* = .002, *d* = 0.46, *BF* = 11.8). Finally, participants needed fewer queries for the high entropy games than for the very high entropy games (*t*(46) = −3.96, *p* < .001, *d* = 0.58, *BF* = 97.2).

**Figure 3:**
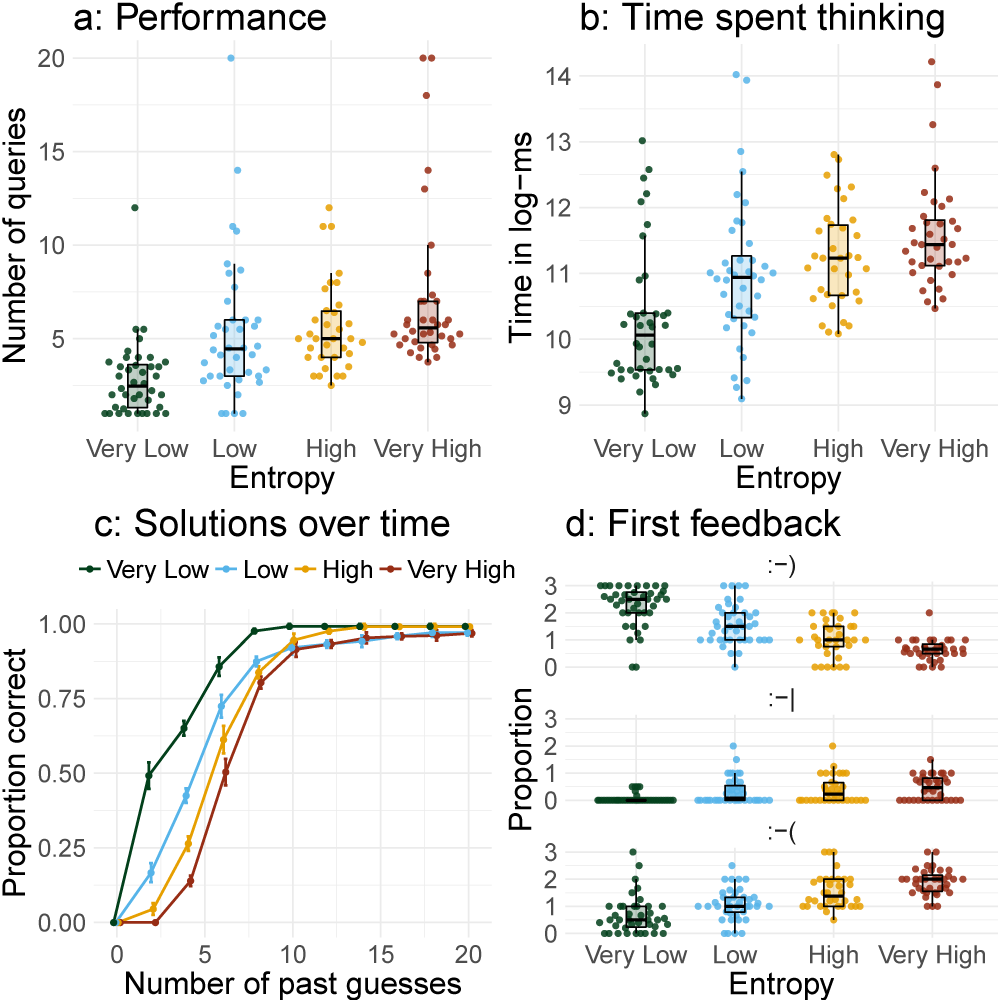
Behavioral results. **a:** Number of queries required to solve a game by entropy condition (ordered from lowest to highest). **b:** Time spent thinking (measured in log-ms per guess) by entropy condition (ordered from lowest to highest). **c:** Proportion of correct guesses in dependency of number of past guesses by entropy condition. **d:** Mean proportional feedback after first guess by entropy condition. Points represent mean per participant. Error bars indicate the standard error of the mean.

Next, we analyzed how much time participants spent thinking to enter a guess by entropy condition (Fig. 3b). Thus, we assessed their mean time to submit a query measured in log-milliseconds. There was a positive average rank-correlation between a game’s entropy and participants’ average time spent thinking, Kendall’s τ = 0.48, *t*(46) = 12.44, *d* = 1.68, *BF* > 100. More specifically, participants spent less time thinking during the very low entropy games than during the low entropy games (*t*(46) = −4.07, *p* < .001, *d* = 0.59, *BF* = 97.2). They also spent less time thinking in the low entropy than in the high entropy games (*t*(46) = −3.68, *p* < .001, *d* = 0.54, *BF* = 45.5). Finally, they spent less time in the high entropy than in the very high entropy games (*t*(46) = −4.05, *p* < .001, *d* = 0.59, *B* > 100).

We also analyzed the proportion of solved games as a function of the number of past guesses, again comparing the different entropy conditions (Fig. 3c). We thus estimated a Bayesian logistic regression of number of past guesses onto the proportion of correct guesses for each of the entropy conditions, using Metropolis-Hastings Markov chain Monte Carlo sampling (implemented in `MCMCpack`, Martin, Quinn, Park, & Park, 2018). The resulting posterior estimate for the effect of number of past guesses onto the probability of guessing correctly was smallest for the very high entropy condition (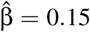, 95%HDI=[0.14, 0.16]). The same estimate was higher for the high entropy condition (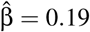, 95%HDI=[0.18, 0.20]), which did not differ meaningfully from the low entropy condition (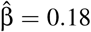, 95%HDI=[0.17, 0.20]). The very low entropy condition showed the highest estimated effect (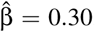, 95%HDI=[0.28, 0.33]). Thus, participants’ solution rates differed meaningfully between entropy conditions, with lower entropy leading to faster rates.

In our last behavioral analysis, we looked at the very first query participants submitted as well as the feedback they received for that query (Fig. 3d). The number of smiling faces received on the very first guess was negatively rank-correlated with entropy condition, τ = −0.51, *t*(41) = −9.80, *p* < .001, *d* = 1.51, *BF* > 100, whereas the number of frowning faces showed a positive rank-correlation, τ = 0.30, *t*(30) = 6.00, *p* <. 001, *d* = 1.06, *BF* > 100. Interestingly, participants adapted their first queries to the entropy condition, leading to a positive rank correlation between the set size of their first query (the number of unique kinds of fruit contained in the query) and the entropy of the generating distribution, τ = 0.40, *t*(46) = 9.00, *p* <. 001, *d* = 1.31, *BF* > 100. Put differently, if the generating distribution was higher in entropy, then participants tested a larger number of different fruits as part of their first query.

### Computational modeling

We now turn to a model-based analysis of participants’ exploration strategies. For this, we first need a formal account of intelligent Mastermind play. Logically, all combinations that are still consistent in round *i* based on the feedback received so far are part of a feasible set 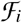. Note that in Entropy Mastermind, not only the feasible codes but also their probabilities (which are not in general equal) are relevant. Code combinations ruled out by prior feedback have zero probability. The remaining items’ probability mass is proportional to the probability of obtaining the item via sampling from the code jar. The effective size of the feasible set is the total number of all non-zero probability codes left in the set. Let the probability that *c*_*i*_ is the hidden code given the current feasible set be denoted *P*(*c*_*i*_). The feasible set is guaranteed to shrink after each round unless a guess *c*_*i*_ is repeated. A general playing strategy consists of (i) identifying the set of feasible combinations 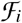 (with 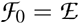), where prior feedback is used to determine which combinations are still viable; and (ii) picking a combination *c*_*i*_ for the next guess. Let us denote the informational usefulness of playing combination *c* in the current round with *u*(*c*), with

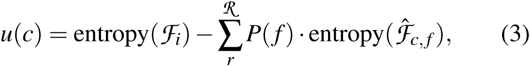

i.e. the difference in entropy (under a particular Sharma Mittal entropy measure with specified order and degree) between the current feasible set and the expected entropy when playing code c. To compute expected entropy, for each possible feedback 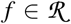, we compute the product of the probability of receiving that feedback *P*(*f*) times the entropy of the updated feasible set 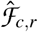 when playing combination *c* and receiving feedback *r*. To compute *P*(*f*) for a given *c*, we look at all the combinations 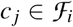, that lead to feedback *f*. To this end, we define a feedback function *h*(*c, c*_*j*_) = *f* that returns the feedback *f* obtained from checking code *c* against code *c*_*j*_. The probability of feedback *f* for code *c* can then be calculated as follows:

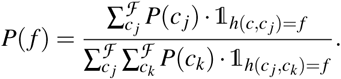

The indicator function 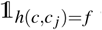 ensures that we only sum over codes *c*_*j*_ that generate the required feedback *f*. The probability of any combination of fruits *c* = *m*_1_*m*_2_…*m*_*n*_ can be computed as

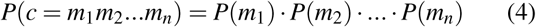

where each *P*(*m*) represents the probability of sampling the corresponding fruit item from the fruit jar. The other term of Equation 3, entropy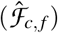, requires us to compute hypothetical feasible sets 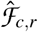. Given the current feasible set 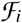, a combination *c* we want to evaluate, and hypothetical feedback *f*, we need to exclude all combinations 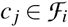 for which *h*(*c, c*_*j*_) ≠ *f*; that is, all combinations *c*_*j*_ that are not consistent with obtaining feedback *f*.

Lastly, one has to assign a utility to a feasible set 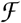. For this, we use the Sharma-Mittal entropy framework to compute the entropy of a probability distribution defined over set 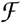, 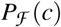. For each combination 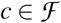

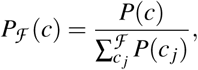

where the nominator *P*(*c*) is computed according to Equation 4 and the denominator is a normalization term.

We assess how well the combination of an entropy-based exploration bonus and the probability of making a correct guess describes players’ guesses over time. For this, we analyzed the last five games of the 34 participants who played at least five games in total. We restricted our analysis to the last five games as our goal was to study strategies used rather than early learning. Next, we calculated the expected information gain for all of the 6 × 6 × 6 possible fruit combinations that a participant could enter on every trial for each participant, given the participant-specific history of queries in a game. We calculated this information gain for every combination of order *r* = [1/16, 1/8, 1/4, 1/2, 1, 2, 4, 8, 16, 32, 64] and degree *t* = [1/16, 1/8, 1/4, 1/2, 1, 2, 4, 8, 16, 32, 64], i.e. 121 models per participant in total. We then combined the probability of a guess being correct with the information gain assessed by the specific entropy measure following Equation 2 to arrive at a value of an action’s usefulness *V* (*a*_*t*_), which we put in a softmax function to calculate choice probabilities:

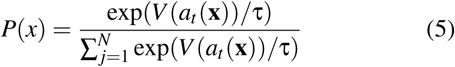

where τ is a free temperature parameter. We followed previous work (Wu et al., 2018; Parpart, Schulz, Speeken-brink, & Love, 2017) and calculated each model’s AIC 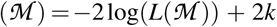 and standardized it using a pseudo-*R*^2^ measure as an indicator for goodness of fit, comparing each model 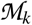 to a random model: 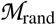, 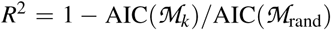.

The results of this analysis revealed a mean pseudo-*R*^2^ of 0.041 over all orders and degrees, which was low but significantly better than chance (*t*(33) = 20.52, *p* < 0.001 *d* = 1.86, *BF* > 100). Moreover, the estimated median temperature parameter was τ = 1.02, indicating a relatively wide spread of predictions. There was a significant negative rank-correlation between the degree parameter and model fit, τ = −0.37, *z* = −5.84, *p* < .001, *BF* > 100, whereas this correlation was not significant for the order parameter, τ = 0.04, *z* = 0.60, *p* = .54, *BF* = 0.3. Thus, even though entropies with smaller degree parameters seemed to generally work better at modeling participants’ queries, there was no meaningful effect of the different order parameters.

The range of pseudo-*R*^2^ values, 0.038 − 0.045, also shows that most of the entropy measures led to similar performance. We also assessed the magnitude of the estimated exploration bonus β (Fig. 4b), which had a mean of 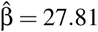, and therefore differed significantly from 0, *t*(33) = 115.47, *p* < .001, *d* = 10.9, *BF* > 100. This means that the final model of participants’ game play had to incorporate both a code’s probability of being correct as well as its potential information gain. Interestingly, areas of the Sharma-Mittal space with higher *r*^2^ also tended to have higher β estimates.

**Figure 4:**
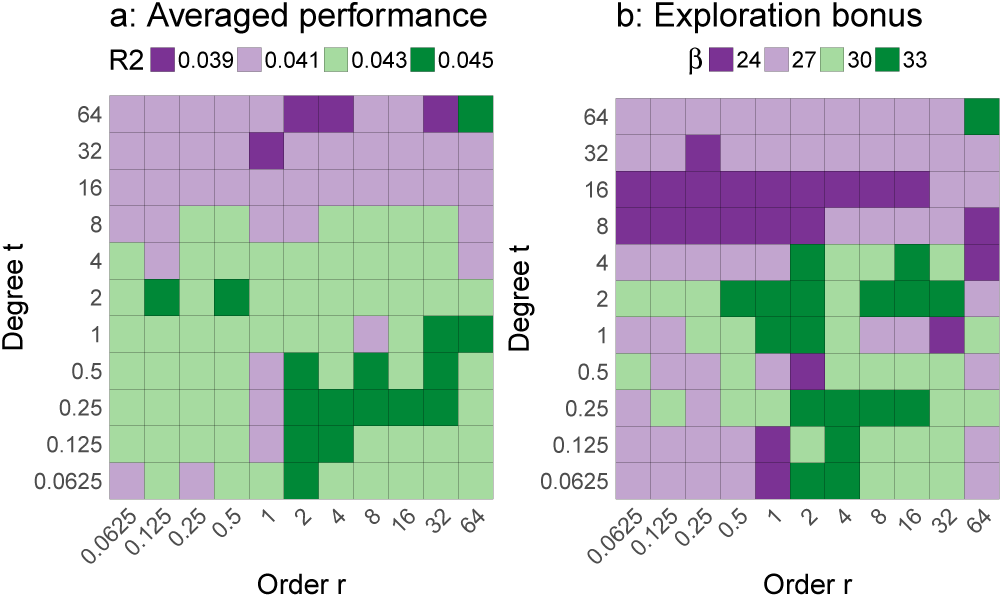
Modeling results. **a:** Averaged *r*^2^ for different Sharma-Mittal parameters. **b:** Estimated exploration bonus β for different Sharma-Mittal parameters.

Finally, we compared how often participants put the most likely fruit into their first query with how often simulated models of different order and degree parameters chose the same fruit in their first query, for each entropy condition (see Fig. 5). The higher degree models chose the most likely fruit more often than people did. Specifically, participants put on average 2.14 of the most likely fruit in their first query in the very low entropy condition, 1.60 in the low entropy condition, 1.26 in the high entropy condition and 0.48 in the very high entropy condition. This analysis therefore corroborated our previous finding that the lower degree entropies better matched participants’ queries. In relation to previous work modeling behavior with the Sharma-Mittal framework, Entropy Mastermind appears to be more similar to experience-based than to description-based probabilistic classification tasks (see Crupi et al., 2018, Fig. 7).

**Figure 5:**
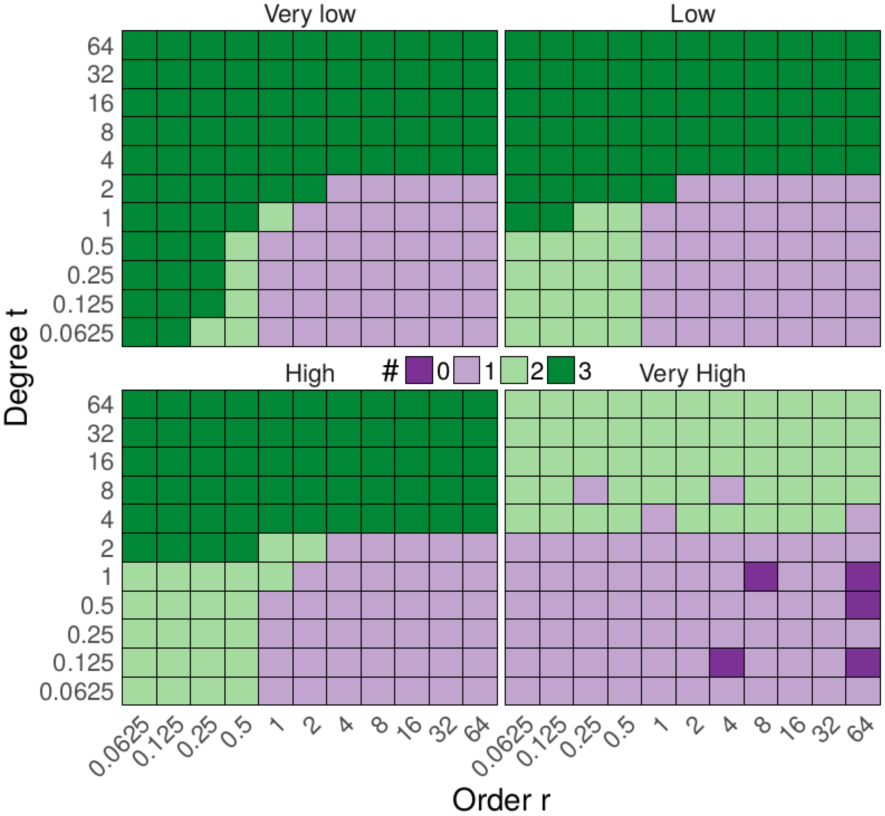
Number of times the most likely fruit was chosen in the first query by simulated entropy models across entropy conditions.

## Discussion and conclusion

We introduced Entropy Mastermind as a game for researching human curiosity and exploration in complex environments. More specifically, we suggest this game as a paradigm for the study of how people select queries to reduce uncertainty under different levels of initial entropy. The complexity of the game resembles aspects of scientific inference (Strom & Barolo, 2011) and life. For instance, in life and in science, it can be a challenge to fully assimilate feedback that we get when we make queries. Entropy Mastermind thus complements existing games, such as Battleship (Gureckis & Markant, 2009), 20-questions (Nelson et al., 2014), or explore-exploit (Wu et al., 2018) tasks.

We found that participants required fewer queries, spent less time thinking about queries and showed faster learning rates if the distribution generating the secret code had lower entropy. They also adapted their queries to the code-generating distribution, and did so in sensible ways. In particular, many of the informational models (Figure 5) used greater proportions of the most-probable fruit in the first guess in lower-entropy conditions; participants also followed this pattern. Thus, one may conclude that people are generally sensitive to different levels of entropy, which is a pre-requisite for a research agenda modeling human exploratory behavior within the Sharma-Mittal space.

Our modeling results paralleled earlier findings from other tasks (Crupi et al., 2018) suggesting that it is easier to identify the value of the degree parameter than of the order parameter in the Sharma-Mittal space. Interestingly, to identify the order parameter a different type of question could be asked, translating higher entropy into difficulty of game play in the sense of the number of queries required to guess the secret code (for the underlying mathematical result see Crupi et al., 2018). Participants could be directly asked which of two code jars would be harder to play Mastermind with (Figure 6).

**Figure 6:**
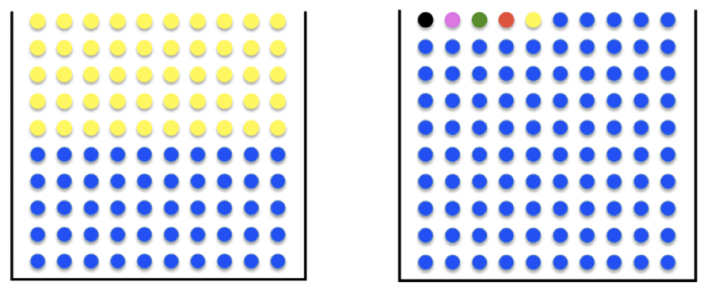
Identifying the order parameter. Which distribution is harder for playing Entropy Mastermind? Shannon entropy (order=1) deems the 50:50 distribution higher entropy, but lower-order entropies deem the 95:1:1:1:1:1 distribution higher entropy.

The general predictive performance of many models was relatively similar and rather low. This might be due to the overall complexity of choices, since there were 216 possible options on every trial, making it difficult to compare among candidate models (also see Parpart et al., 2017).

The difficulty of modeling could also be due to participants using cognitive shortcuts, as has been observed in other domains of active learning (Bramley et al., 2017). Furthermore, it is unlikely that participants evaluate the usefulness of all possible queries at each time point. Instead, they might approximate a query’s usefulness by sampling and reusing past hypotheses, as has been shown in other domains of human reasoning and hypothesis evaluation (Dasgupta, Schulz, & Gershman, 2017; Dasgupta, Schulz, Goodman, & Gershman, 2018; Lieder, Griffiths, & Hsu, 2018). Future studies should therefore investigate both heuristic strategies (Gigerenzer & Gaissmaier, 2011) and boundedly rational approaches (Griffiths, Lieder, & Goodman, 2015). Adaptive experimental designs (Cavagnaro, Myung, Pitt, & Kujala, 2010) could also be used to maximally discriminate among models.

Summing up, we propose Entropy Mastermind as a promising paradigm for investigating human exploration behavior in complex hypothesis testing scenarios. In related research we are assessing whether Entropy Mastermind can be used as an educational tool for primary or secondary school students, and for studying the effects of emotional states on strategies used and information search efficiency. Although our current modeling framework did not fully map out the space of exploration behavior, we believe that combining the Sharma-Mittal space of entropy measures with an enjoyable game rich in scientific history can further inform our theories of self-directed learning. We will keep exploring.

## Acknowledgments

ES is supported by the Harvard Data Science Initiative. This work was supported by grant NE 1713/2 to JDN from the Deutsche Forschungsgemeinschaft as part of the priority program New Frameworks of Rationality (SPP 1516). We thank Eloisa Bentivegna, Neil Bramley, Vincenzo Crupi, Florian Ellsaesser, Alberto Feduzi, Flavia Filimon, George Kachergis, Laura Martignon, Bjoern Meder, Elif Oezel, Anselm Rothe, Azzurra Ruggeri, Katya Tentori, John Wong and the iSearch research group for helpful comments and ideas.

## Footnotes

1 Note that the parts of Eq. 1 are additive. Thus, even a query that has a probability of 0 of being the true code can still be chosen if it offers enough informational value.

